# Intronic miR342 is a master regulator of cellular glycolysis in Foxp3^+^ regulatory CD4 T cells

**DOI:** 10.1101/2023.09.18.558294

**Authors:** Rongzhen Yu, Silei Li, Sohee Kim, Giha Kim, Li-Fan Lu, Dongkyun Kim, Booki Min

## Abstract

Foxp3^+^ regulatory T (Treg) cells are a subset of CD4 T cells that play a potent and indispensable role in regulating immunity and tolerance. The precise mechanisms by which Treg cells mediate such functions have extensively been explored, and there are many cellular and molecular factors that are instrumental for adequate Treg cell functions. miRNAs, small non-coding RNA molecules, are one of the factors capable of controlling Treg cell functions. In this study, we report that miR-342 is essential for Treg cells to mitigate autoimmune inflammation in the central nervous system. Utilizing novel mouse models with Treg cell-specific miR-342 deficiency or overexpression, we demonstrate that miR-342 expression in Treg cells, while dispensable for immune homeostasis at steady-state conditions, is necessary for Treg cells to control inflammatory responses. Mechanistically, we found that Treg cells deficient in miR-342 display dysregulated metabolic profiles, elevated glycolysis and decreased oxidative phosphorylation, a metabolic phenotype associated with functionally defective Treg cells. Interestingly, miR-342-dependent metabolic dysregulation was observed in Treg but not in Th1 type cells. In support, miR-342-mediated Rictor targeting was found in Treg but not in Th1 type cells. Collectively, our findings uncover that miR-342 may serve a master regulator specific for metabolism and functions in Treg cells.

## Introduction

Foxp3^+^ regulatory T (Treg) cells play indispensable roles in the immune system. Defective Treg cell function or generation could result in uncontrolled systemic or tissue specific immune activation, while intensified Treg cell function could interfere with protective immunity ^1^. The former example includes chronic autoimmune inflammation, where adequate Treg cell functions are dwindled. In the latter case such as chronic pathogen infection or tumor, Treg cells acquire better suppressive functions that inhibit effector immunity, enabling certain infectious agents to evade the host immune responses or preventing cytotoxic T cells from killing tumor cells ^2, 3^. The mechanisms involved in variable Treg cell functions are not well understood, and identifying the key cellular and molecular pathways harnessing Treg cell function is a subject of ongoing investigation.

microRNA (miRNA) is a ∼22 nucleotide non-coding RNA that plays a critical role in gene regulation through binding the 3’ untranslated region (UTR) of target genes ^4^. It is estimated that miRNAs may modulate ∼60% of protein encoding genes in the human genome ^5^. miRNAs also play a crucial role in Treg cell development and functions. Dicer, a cytoplasmic RNase III mediating pre-miRNA processing, has previously been shown to be critical for Treg cells based on the findings that Dicer deletion in Treg cells cause rapid fatal autoimmune inflammation ^6^. Some miRNAs including miR-146 and miR-155 are highly expressed in Treg cells, regulating Treg cell suppressive functions and proliferative capacity, respectively ^7–9^. We previously demonstrated that miR-342 plays a role in Treg cell suppressive function by modulating cellular metabolic profiles, in part targeting metabolic regulator, Rictor ^10^.

miR-342 is an intronic miRNA located within the third intron of the host gene ENAH-VASP-like (Evl). miR-342 does not have its own regulatory element, and its expression is believed to be controlled by the host gene ^11^. miR-342 plays multiple roles in various cell types and cancer cells. For example, it is known to regulate Myc transcriptional activity via direct repression of E2F1 transcription factor in human lung cancer cells ^12^. miR-342-3p also suppresses hepatocellular carcinoma proliferation through inhibiting IGF1R mediated Warburg effect, suppressing IGF1R-mediated PI3K/Akt/Glut1 signaling pathway and dampening glycolysis by decreasing glucose uptake, lactate generation, and ATP production ^13^.

Cellular metabolism is considered instrumental for adequate immune functions. Distinct contribution of metabolic pathways in Treg cell biology has extensively been examined. In general, Treg cells prefer oxidative glycolysis, a process in which glucose is broken down to pyruvate, which then enters the TCA cycle, instead of anaerobic glycolysis in which pyruvate is converted to lactate ^14^. Glycolysis, while necessary for Treg cell proliferation, is known to impair Treg cell suppressive capacity ^15^. We previously reported that miR342 inhibition in Treg cells using an antagomir diminishes optimal Treg cell suppressive function to limit inflammatory responses ^10^. On the other hand, miR-342 mimic treatment of Treg cells improves the function in vivo ^10^.

In this study, we utilized newly generated Treg cell-specific miR-342 deficient and overexpression animal models and investigated the role of miR-342 in Treg cell function. We found that miR-342 deficiency in Treg cells greatly impairs the suppressive ability to dampen autoimmune inflammation. Conversely, overexpression of miR-342 in Treg cells was able to enhance Treg cell suppressive function. RNAseq analysis uncovered that metabolic processes are the pathway highly enriched by miR-342 deficient Treg cells. Specifically, ECAR was drastically increased in miR-342 deficient Treg cells, and OCR, an indication of oxidative phosphorylation, was significantly reduced by miR-342 deficiency. Interestingly, miR-342-dependent metabolic regulation was observed in Treg but not in Th1 type effector cells, suggesting a cell type specific role of miR-342. Finally, we compared the expression of previously reported targets regulated by miR-342 and found that Rictor expression was significantly affected by miR-342 in Treg while not in Th1 type cells, suggesting a unique function of miR-342 in Treg lineage cells.

## Materials and Methods

### Animals

Foxp3^YFPCre^ (B6.129(Cg)-Foxp3tm4(YFP/icre)Ayr/J, #016959) and Foxp3^DTR^ (B6.129(Cg)-Foxp3^tm3^(Hbegf/GFP)^Ayr/J^, #016958) mice were purchased from the Jackson Laboratory (Bar Harbor, ME). The generation of miR-342-floxed mice was commissioned by Biocytogen (Beijing, China). The targeting construct for miR-342 overexpressing Tg mice was generated similar to what was described previously ^16^. All the mouse protocols were approved by the Institutional Animal Care and Use Committee (IACUC) of the Northwestern University. The mice were maintained under specific pathogen-free conditions at the Northwestern University Feinberg School of Medicine.

### Experimental Autoimmune Encephalomyelitis (EAE) induction

Mice were immunized subcutaneously into the rear flanks with 1:1 emulsion of MOG_35-55_ peptide (300μg, BioSynthesis, Lewisville, TX) in complete Freund’s adjuvant (CFA) containing 5 mg/ml supplemented Mycobacterium tuberculosis H37Ra (Difco, Detroit, MI). Mice were injected with 200 ng pertussis toxin (Sigma, St. Louis, MO) intraperitoneally on day 0 and 2 post immunization. EAE development was monitored and graded on a scale of 0 to 5: 0, no symptoms; 1, flaccid tail; 2, hind limb weakness; 3, hind limb Paralysis; 4, fore limb weakness with hind limb paralysis; and 5, moribund or death. For passive EAE induction, Foxp3^DTR^ mice were induced for EAE. At disease onset (typically day 7 post immunization), mice were injected with 1μg Diphtheria toxin (Sigma) and adoptively transferred with 2 x 10^6^ in vitro generated iTreg cells.

### Flow cytometry

Single cell suspensions were prepared from lymph nodes (LN), brain and spinal cord as previously described ^17^. The following Ab were used: anti-CD4 (RM4–5), anti-CD8 (53-6.7), anti-CD44 (IM7), anti-CD25 (PC61.5), anti-GITR (DTA-1), anti-ICOS (C398.4A), anti-Lag3 (C9B7W), anti-Foxp3 (FJK-16s), anti-PD1 (J43), anti-GITR (DTA-1), and anti-Nrp1 (3DS304M). For intracellular cytokine detection, cells were stimulated for 4 h with

PMA (10ng/mL, Millipore-Sigma) and ionomycin (1μM, Millipore-Sigma) for 4 h in the presence of 2μM monensin (Calbiochem, San Diego, CA) during the last 2 h of stimulation. Cells were fixed with 4% paraformaldehyde, permeabilized, and stained with the following Ab: anti-IL-17 (TC11-18H10), anti-IFNγ (XMG1.2), anti-TNFα (TN3-19), anti-GM-CSF (MP1-22E9). The Abs were purchased from eBioscience (San Diego, CA), BD Biosciences (San Diego, CA) and BioLegend (San Diego, CA). Stained cells were acquired with a FACS Symphony A1 (BD Biosciences) and analyzed using FlowJo software (TreeStar, Ashland, OR).

### In vitro differentiation

FACS sorted naive CD4 T cells were stimulated with immobilized anti-CD3 (2C11) plus anti-CD28 (37.51) mAbs for 3 days. For Th1 differentiation, IL-12 (Peprotech, Rocky Hill, NJ, 10ng/ml) and IL-2 (20U/ml) were added. For iTreg cell differentiation, rTGFb1 (Peprotech, 5ng/ml) and rIL-2 (100U/ml) were added in the culture.

### Real time quantitative PCR

Mice with EAE were euthanized and perfused with PBS. The brain and spinal cords were isolated and total RNA was extracted using a TRIzol reagent according to the manufacturer’s instructions (Invitrogen). cDNA was then generated using a MMLV reverse transcriptase (Promega, Madison, WI). qPCR analysis was performed using a QuantStudio 3 Real-Time PCR System (Applied Biosystems, Waltham, MA) using a Radiant qPCR mastermix (Alkali Scientific, Fort Lauderdale, FL) or SYBR green mastermix (Applied Biosystems). The data were normalized by housekeeping *Gapdh* gene and then compared to the control group. Taqman primers used in this study are: pri-miR-342 (Mm03307009_pri), pre-miR-342 (Mm04238212_s1), miR-342 (002260), Evl exon3-4 (Mm00468401_m1), Evl exon7-8 (Mm00468405_m1), Foxp3 (Mm00475162_m1), Hk2 (Mm00443385_m1), Ldha (Mm01612132_g1), and Pgk1 (Mm00435617_m1).

### RNAseq

RNA was isolated from Tregs isolated from the CNS tissues using a RNeasy Micro kit (Qiagen). The quality of all samples was assessed on a Fragment Analyzer electrophoresis system (Agilent). Total RNA was normalized prior to oligo-dT capture and cDNA synthesis with SMART-seq v4 kit (Takara). The resulting cDNA was quantitated using a Qubit 3.0 fluorometer. Libraries were generated using the Nextera XT DNA library prep kit (Illumina). 25 million reads per sample was performed using a HiSeq2500 (Illumina) on a Rapid run flow cell using a 100 base pairs, paired end run. FASTQ files were uploaded and analyzed on Galaxy Project (https://usegalaxy.org). Read qualities were examined by FastQC module (http://www.bioinformatics.babraham.ac.uk/projects/fastqc). RNA sequencing reads were aligned to the genome via STAR version 2.7.8a (https://doi.org/10.1093/bioinformatics/bts635) and the GRCm38.p4(M10) mouse genome annotation file was used as a reference ^18^. The following parameters were used: pair-ended, mm10 reference genome, 150 bp length of the genomic sequence around annotated junctions, and all other default parameters. Aligned reads were counted using FeatureCount ^19^ with enabled multi-mapping, multi-overlapping features, and assigned fractions. Counted genes were further normalized by DESeq2 for differential expression analysis ^20^ and results were visualized by ggplot2 in Rstudio ^21^. GSEA was performed using GSEA software from UC San Diego and Broad Institute ^22, 23^. Differentially expressed genes were also subjected to Metascape analysis ^24^. Top-level gene ontology biological processes were examined. Raw and processed sequencing data have been deposited at NCBI Gene Expression Omnibus under accession umber GEO:GSE241986.

### Seahorse flux assay

Glycolysis stress test kit and Seahorse XFe96 (Agilent, Santa Clara, CA) were used to evaluate oxidative phosphorylation and glycolysis. Briefly, XF 96-well plates were coated using Cell-Tak (BD Biosciences) according to the Seahorse protocol, and cells were plated at a concentration of 3 x 10^5^ cells per 50 µl of XF assay medium containing 2 mM glutamine. After 30 min incubation in CO^2^-free incubator at 37°C, cells were spun down to enhance cell adherence, and an additional 100 µl of XF assay medium was added per well. Oxygen consumption rate (OCR) was measured by sequentially injecting oligomycin (1μM), FCCP (2μM), and rotenone/antimycin A (0.5μM). Extracellular acidification rates (ECAR) were measured by sequentially injecting glucose (1mM), oligomycin (1μM), and 2-deoxyglucosee (2-DG, 2μM). Basal and Max OCR and ECAR were calculated according to the manufacturer’s instructions.

### Statistics

Statistical significance was determined by the Mann-Whitney test using a Prism software (GraphPad, San Diego, CA). p<0.05 was considered statistically significant.

## Results

### Generation of Treg cell-specific miR-342 deficient mice

To investigate the role of miR-342 in Foxp3^+^ Treg cells we generated miR-342-floxed mice and then crossed to Foxp3^CreYFP^ transgenic mice (Fig 1A). To validate Treg cell-specific miR-342 deletion, conventional CD4 and Treg cells were FACS sorted from Foxp3^Cre+^ mir342^fl/fl^ and control Foxp3^Cre-^ mir342^fl/fl^ mice. Treg cell-specific deletion was verified by genomic DNA PCR analysis. As shown in Fig 1B, 5’F and 3’R primers amplified miR-342-deleted allele only in tTreg cells from Foxp3^Cre+^ mice. On the other hand, miR-342-containing allele remained intact in tTreg cells from Foxp3^Cre-^ control mice or in conventional CD4 T cells in both strains. Therefore, Treg cell-specific miR-342 deletion is confirmed.

**Figure 1.**
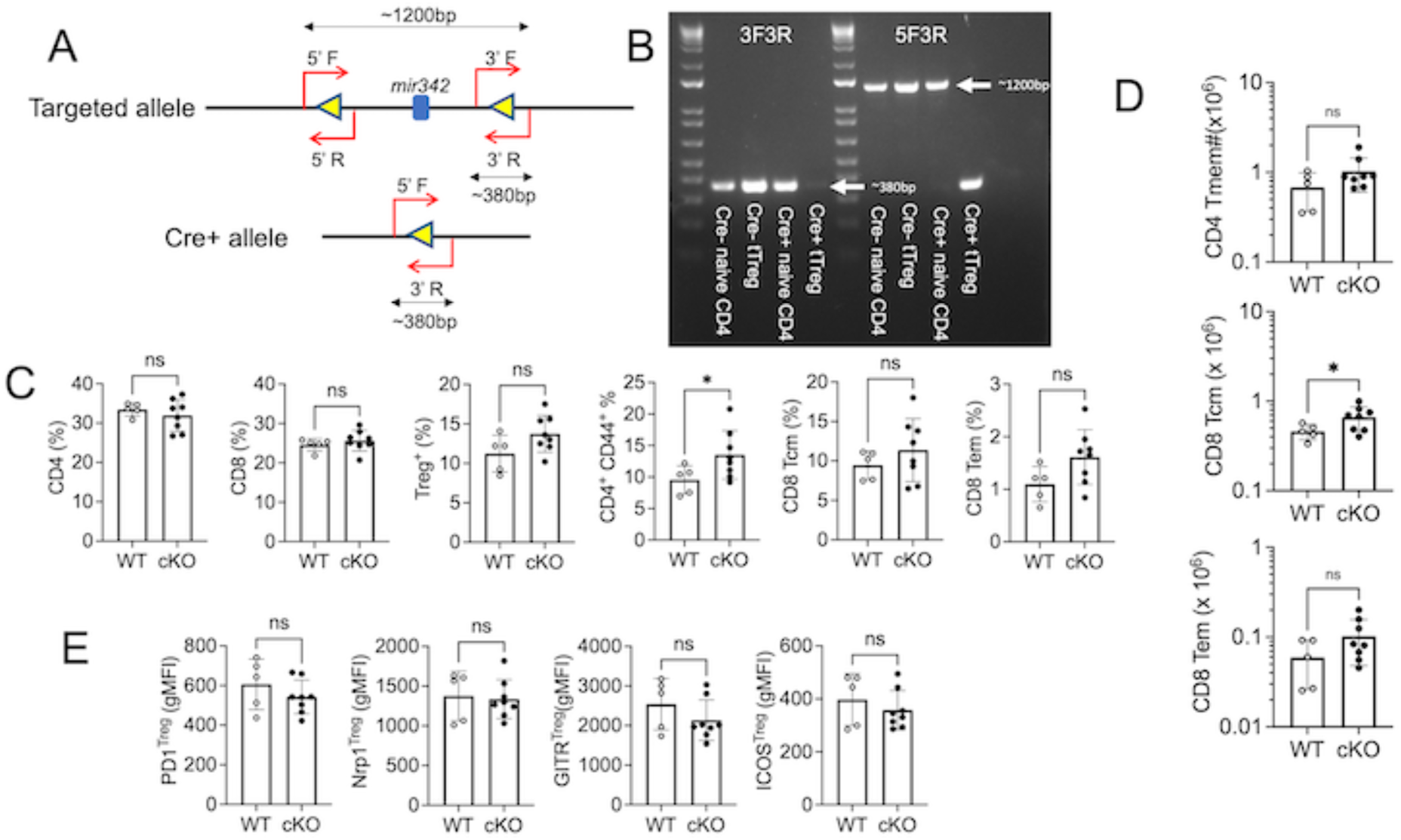
Treg cell-specific miR-342 mutant mice. (A) Targeting strategy. Loxp sites were introduced to the mir342 locus. Genotyping primers and expected size of unfloxed and floxed alleles are shown. (B) PCR analysis of gDNA using the primers, 5’F, 3’F, and 3’R. Naive and CD25^+^ Treg cells were FACS sorted from Foxp3^Cre+^ miR342^fl/fl^ and Foxp3^Cre-^ miR342^fl/fl^ mice. Genomic DNA PCR analysis was performed. (C) Lymph nodes of Treg^Δ*mir*342^ and Treg^WT^ mice were examined for CD4, CD8, and Treg cells. (D) Absolute numbers of CD4 and CD8 memory phenotype cells were enumerated. (E) Treg cell surface markers were examined by flow analysis. Each symbol represents individually tested mouse. *, p<0.05.

Treg cell-specific miR-342-deficient mice (referred to as Treg^τι*mir*342^ hereafter) were viable and did not display any adverse signs of immune activation in the primary and secondary lymphoid tissues. Total numbers of lymph node cells were comparable between the mutant and control mice (Supp Fig S1). Proportions of CD4, Treg, and CD8 T cells were also similar between the groups (Fig 1C). While CD44^high^ memory phenotype CD4 T cells were found significantly increased in Treg^τι*mir*342^ mice, their absolute numbers remained comparable (Fig 1C and 1D). Similarly, the frequency of central and effector memory phenotype CD8 T cells were slightly increased in Treg^τι*mir*342^ mice, although it did not reach statistical significance (Fig 1C). Yet, the absolute number of central memory CD8 T cells was found greater in Treg^τι*mir*342^ mice (Fig 1D). However, we found that Treg cell associated markers including PD1, Nrp1, GITR, and ICOS were expressed at comparable levels (Fig 1E). Collectively, these results demonstrate that miR-342 deficiency in Treg cells does not have a major impact on peripheral T cell homeostasis under steady-state conditions.

*Treg cell-specific miR-342 deficient mice are susceptible to autoimmune inflammation* To investigate the role of miR-342 in Treg cell function during inflammation we utilized experimental autoimmune encephalomyelitis (EAE), a myelin antigen-specific autoimmune inflammation in the central nervous system, as a model system. As shown in Fig 2A, Treg^τι*mir*342^ mice developed exacerbated disease compared to wild type control mice. Supporting the severe disease are immune profiles of the target CNS tissues. CD4 T cells infiltrating the CNS were markedly increased in Treg^τι*mir*342^ mice (Fig 2B). CD4 T cells displaying proinflammatory cytokine signatures, namely IL-17, TNFα, IFNγ, and GM-CSF, were more abundant in Treg^τι*mir*342^ mice (Fig 2C). CNS infiltrating Treg cells were also found increased in the absence of miR-342 in Treg cells (Fig 2B). Treg cells from Treg^τι*mir*342^ mice expressed comparable levels of ICOS and CD25, while their expression of GITR and Lag3 was significantly reduced by miR-342 deficiency (Fig 2D). Treg cell-intrinsic roles of miR-342 was further examined using miR-342 deficient iTregs. To this end, we generated miR-342 deficient iTreg cells by stimulating naive CD4 T cells isolated from CD4^Cre^ miR342^fl/fl^ (T^τι*mir*342^) mice. Of note, T cell specific miR-342 deficient (T^τι*mir*342^) mice did not show any signs of immune activation, suggesting that miR-342 expression in T cells is dispensable for the development and homeostasis at steady state conditions (Supp Fig S2). In vitro generation of Foxp3^+^ Treg cells was not affected by miR-342 deficiency as we see similar frequency of Foxp3^+^ cells (data not shown). Likewise, Foxp3 mRNA expression of iTreg cells was comparable between wild type and miR-342 deficient Treg cells (Fig 2E). Mature- and pre-miR-342 expression was drastically reduced in iTreg cells generated from T^τι*mir*342^ mice, validating the lack of miR-342 (Fig 2E). We then conducted in vivo Treg suppression assay by transferring iTreg cells into Foxp3^DTR^ mice that were immunized with MOG peptide for EAE induction and injected with DTx at the disease onset (typically day 7 post induction) to deplete endogenous Treg cells. Wild type iTreg cell recipients developed a typical EAE with mild severity and partial remission after the initial peak of the disease (Fig 2F). On the other hand, miR-342 deficient iTreg cell recipients developed severe EAE with no signs of remission, suggesting impaired Treg cells’ ability to mitigate the disease (Fig 2F). In support of the disease severity, the number of CNS infiltrating CD4 T cells from miR-342 deficient iTreg recipients was significantly higher compared to those from control iTreg recipients (Fig 2G). Therefore, these results demonstrate miR-342 dependent Treg cell functions to control autoimmunity.

**Figure 2.**
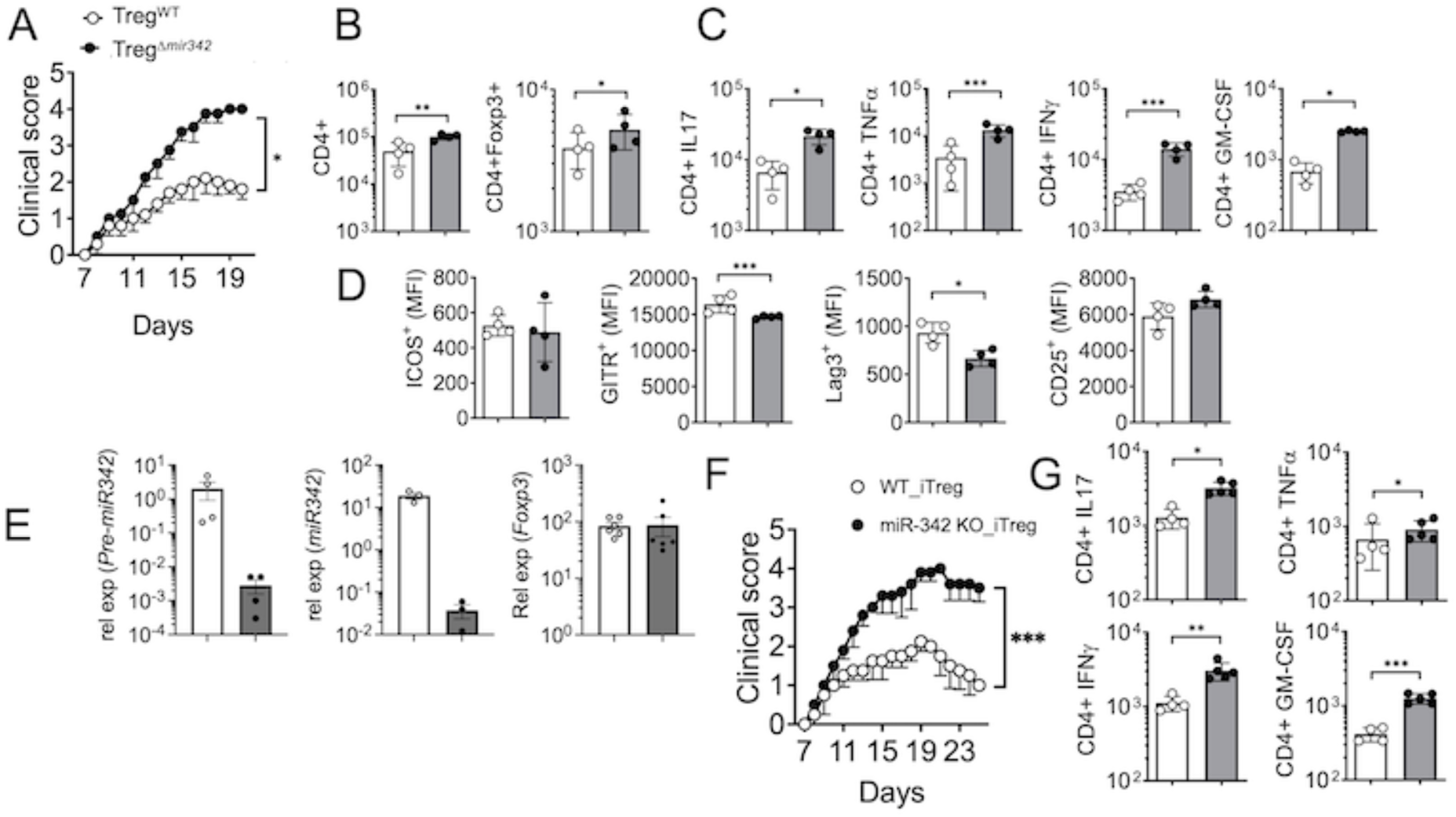
miR-342 deficient Treg cells are unable to control autoimmune inflammation. (A) Groups of Treg^Δ*mir*342^ and Treg^WT^ mice (n=5) were induced for EAE. Clinical score was daily monitored. (B and C) The mice were sacrificed at the peak of the disease. CNS infiltrating CD4 T and Treg cells were examined. (D) Treg cell expression of the indicated surface molecules was measured by flow cytometry. (E) Naive CD4 T cells from T^Δ*mir*342^ and wild type control mice were in vitro stimulated under iTreg cell condition. Foxp3, pre-miR-342, and miR-342 expression was determined by qPCR analysis. (F) In vitro generated iTreg cells were adoptively transferred into Foxp3^DTR^ mice that were induced for EAE and injected with DTx for endogenous Treg cell depletion. EAE development was monitored. The experiments were repeated twice and similar results were obtained.

### miR-342 deficient Treg cells exhibit dysregulated metabolic activity

To investigate the mechanisms underlying miR-342 dependent Treg cell functions, we conducted a RNAseq experiment using wild type and miR-342 deficient iTreg cells. We found that 89 and 60 genes were significantly upregulated in miR-342 deficient and wild type iTreg cells, respectively (Fig 3A). In particular, genes upregulated in miR-342 deficient iTreg cells included genes associated with cellular metabolism, such as DNA damage inducible transcript 4 (Ddit4), Slc16a3, Ldha, and Hk2. Ddit4 is induced under stress responses and known to downregulate mTOR activity ^25^, while Slc16a3, also known as Monocarboxylate transporter 4 (MCT4), exports lactate in cancer cells ^26^. Ldha (lactate dehydrogenase A) converts pyruvate to lactate during glycolysis ^27^, and Hexokinase 2 (Hk2) phosphorylates glucose to produce glycose-6-phosphate, a key rate-limiting step during aerobic glycolysis pathway ^28^. Metascape analysis from significantly upregulated gene list uncovered that metabolic process was the top pathways enriched in these cells, suggesting that miR-342 may regulate cellular metabolic processes (Fig 3B). Gene Set Enrichment Analysis (GSEA) further demonstrated that metabolic pathways enriched in miR-342 deficient Treg cells include metabolic pathways of sugars (glycolysis/gluconeogenesis, galactose metabolism, and amino sugar and nucleotide sugar) as well as of amino acids (metabolisms related to alanine aspartate and glutamate, arginine and proline, glycine serine and threonine) (Fig 3C). Especially interesting is that every gene encoding enzymes in cellular glycolysis were found increased by miR-342 deficiency (Fig 3D). Moreover, pathways of mTORC1 signaling, glycolysis, hypoxia, and inflammatory responses were highly enriched in miR-342 deficient Treg cells (Supp Fig S3). Therefore, miR-342 deficient Treg cells may express dysregulated metabolic profiles, especially glycolysis.

**Figure 3.**
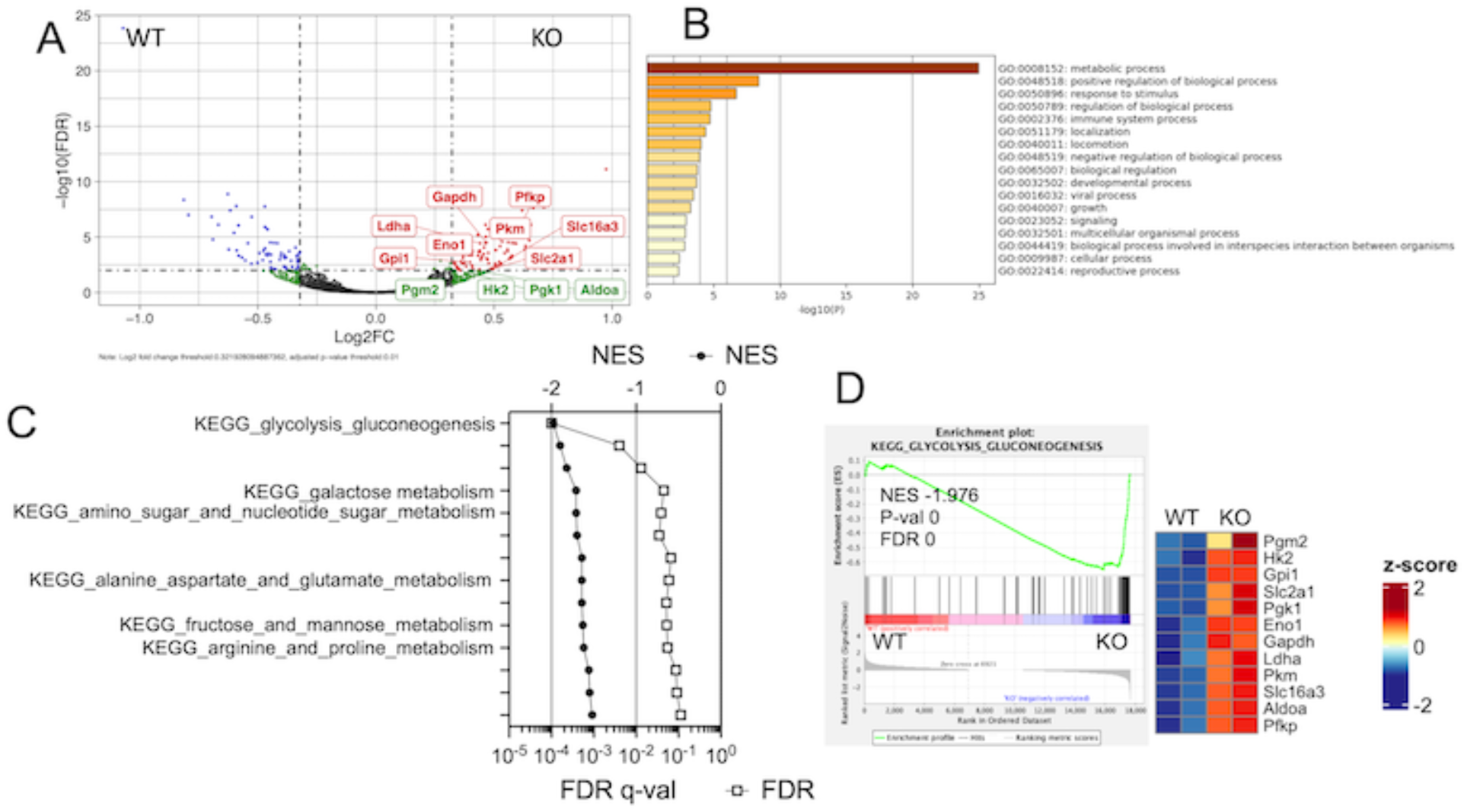
Transcriptomic analysis of miR-342 deficient Treg cells. (A) In vitro generated iTreg cells were used for RNAseq analysis. A volcano plot demonstrating differentially expressed genes in wild type and miR-342 deficient Treg cells. (B) Metascape analysis was performed using genes upregulated in miR-342 deficient Treg cells. (C) Gene set enrichment analysis was performed. Gene sets highly enriched in miR-342 deficient Treg cells were ploted for its normalized enrichment score (NES) and FDR q-val. (D) GSEA plot for the KEGG pathways for glycolysis and gluconeogenesis is shown. Heatmap plot shows genes of glycolytic pathways.

### miR-342 deficient Treg cells display altered cellular metabolism

We previously reported using both tTreg and iTreg cells that Treg cells treated with miR-342 antagomir exhibit increased glycolysis and decreased oxidative phosphorylation and that miR-342 mimic treated Treg cells display opposite metabolic profiles, i.e., decreased extracellular acidification rate (ECAR) and increased oxygen consumption rate (OCR), suggesting that miR-342 may play a role in cellular metabolism ^10^. miR-342 was also previously reported to control Myc transcriptional activity via direct repression of E2F1, a key transcription factor regulating cell cycle progression as well as cellular metabolism, in lung cancer cells ^12^. To further investigate the direct role of miR-342 in Treg cell metabolism, we carried out a Seahorse experiment measuring ECAR and OCR to measure glycolysis and oxidative phosphorylation, respectively, from miR-342 deficient iTreg cells. As shown in Fig 4A, ECAR was substantially increased in miR-342 deficient Treg cells, consistent with the previous findings ^10^. To our surprise, miR-342 deficient Th1 type effector cells, a glycolytic cell type displaying high ECAR, showed only marginal, albeit statistically significant, increase in their ECAR (Fig 4B). On the other hand, miR-342 deficient Tregs displayed significantly diminished OCR, supporting the previous finding that miR-342 supports oxidative phosphorylation of Tregs (Fig 4C). However, miR-342 deficiency in Th1 type effector cells resulted in opposite, small but statistically significant, effects on oxidative phosphorylation (Fig 4D). Therefore, these results suggest that miR-342 has impact on cellular metabolism to support oxidative phosphorylation and that the impact appears to be a cell type specific, especially in Treg lineage cells.

**Figure 4.**
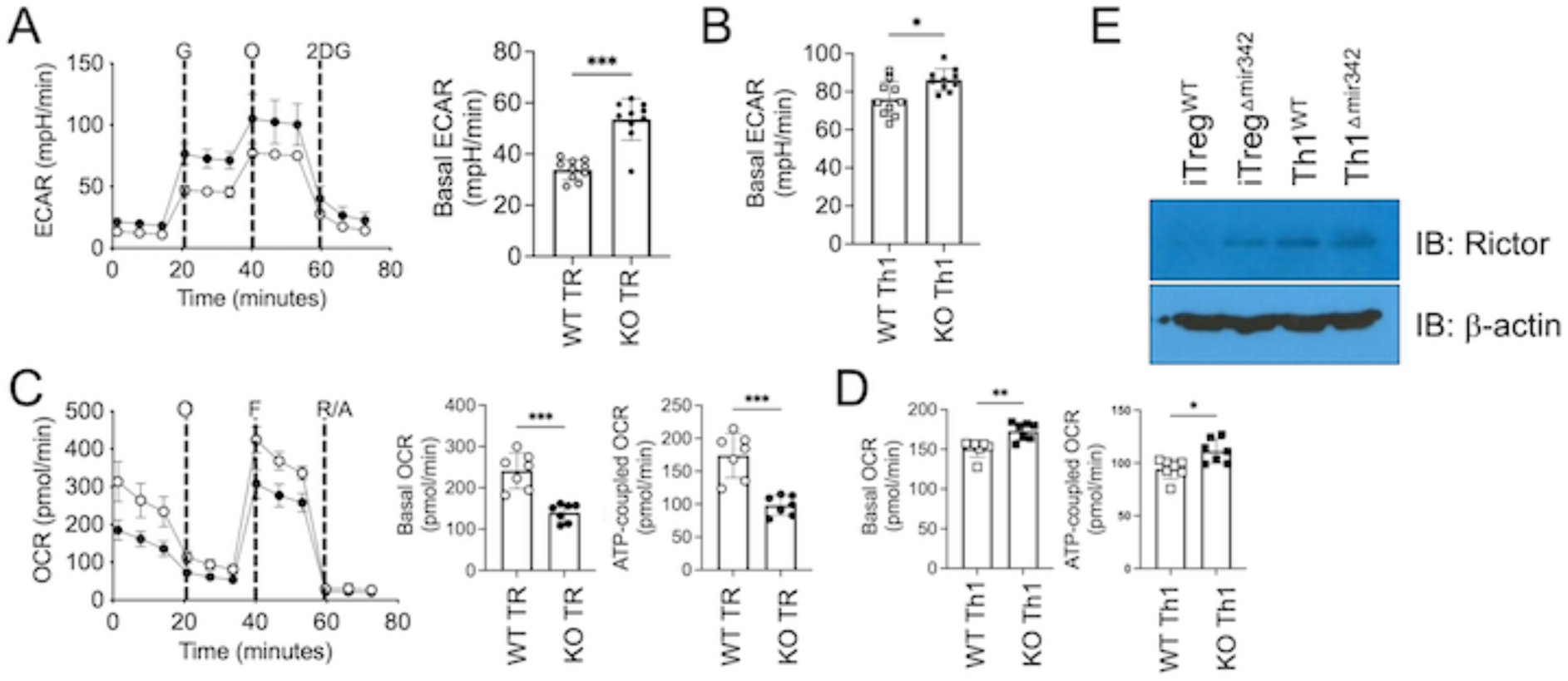
Seahorse analysis of miR-342 deficient and wild type control Treg and Th1 type effector cells. In vitro generated miR-342 deficient and wild type iTreg and Th1 effector CD4 T cells were subjected to Seahorse assay. ECAR (A and B) and OCR (C and D) were measured. (E) Western blot analysis to detect Rictor expression in wild type or miR-342-deficient Treg and Th1 type cells. *, p<0.05; **, p<0.01; ***, p<0.001.

miR-342 is known to regulate Myc activity by controlling E2F1, a Myc-cooperating transcription factor, in cancer cells ^12^. However, we found no differences in E2F1 or Myc expression in Treg cells deficient in miR-342 (data not shown). We previously demonstrated that Rictor, a key component of Rapamycin-insensitive mTORC2 complex, could potentially be targeted by miR-342 in Treg cells and modulate cellular metabolic activity ^10^. Indeed, we observed higher Rictor expression in miR-342 deficient Treg cells (Fig 4E). In agreement with relatively minor impact of miR-342 on Th1 type effector cell metabolism (Fig 4B and 4D), we found that miR-342 deficiency did not affect Rictor expression in Th1 type cells (Fig 4E). Therefore, miR-342-dependent regulation of cellular metabolism and of Rictor expression appears to be unique to Foxp3^+^ Treg cells.

### miR-342 overexpression in Treg cells mitigates EAE development

To further test the role of miR-342 in Treg cell function, we next utilized miR-342 overexpressing Treg cells to test its impact on Treg cell function. Treg cell-specific miR-342 overexpression (miR-342 Tg) mice were generated as previously reported ^16^. miR-342 Tg mice were healthy and viable. Treg cells in the secondary lymphoid tissues were comparable, suggesting that miR-342 overexpression does not affect Treg cell generation (Fig 5A). Proportion of conventional T cells expressing effector/memory phenotypes was also comparable (Fig 5A). Moreover, Treg cell phenotypes measured by proliferation, GITR, Nrp1, and Foxp3 expression were also similar regardless of miR-342 overexpression (Fig 5B). Therefore, miR-342 overexpression in Treg cells does not appear to affect Treg cell development and immune homeostasis in the periphery. To test the effect of miR-342 expression in Treg cell function in vivo, in vivo suppression experiment using iTreg cells was chosen as above. First, naive CD4 T cells from miR-342 Tg mice were stimulated under Treg cell differentiation condition. Foxp3 mRNA expression was similar between miR-342 Tg and control iTreg cells (Fig 5C). Pre- and mature miR-342 expression was found greater in Tg iTreg cells as expected (Fig 5C).

**Figure 5.**
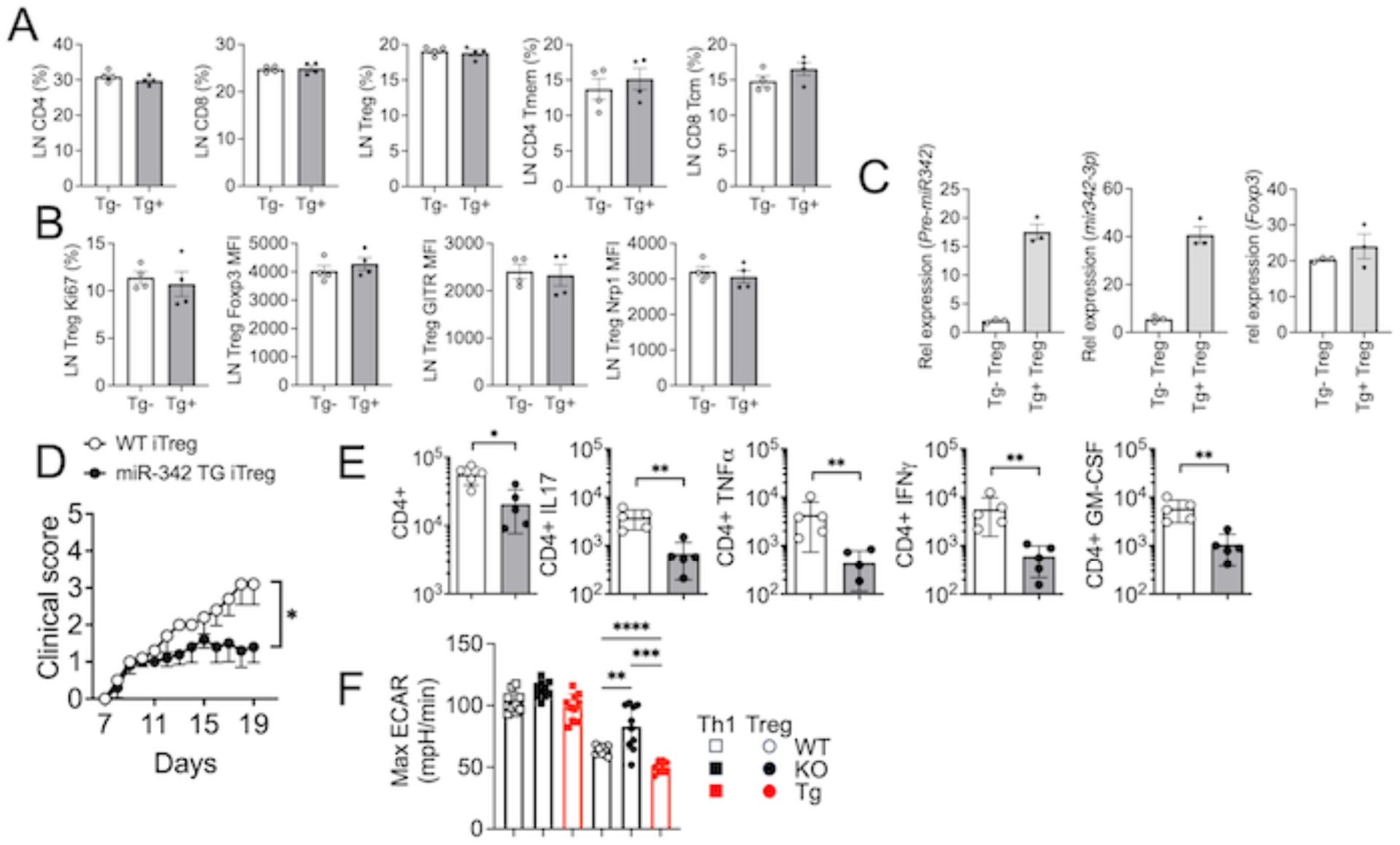
miR-342 overexpression mice are resistant to EAE. (A) Treg cell-specific miR-342 overexpression (Tg) mice were examined. Shown are the proportions of lymph node CD4, CD8, and Treg cells. (B) Treg cell proliferation and surface phenotypes were measured by flow analysis. (C) iTreg cells were generated from miR-342 Tg and wild type control mice. Pre-miR-342, miR-342, and Foxp3 expression was determined by qPCR analysis. (D) Adoptive transfer of miR-342 Tg and wild type iTreg cells was performed as described above. (E) CNS infiltrating CD4 T cells from the aforementioned groups were examined at the peak of the disease. (F) iTreg and Th1 type effector CD4 T cells from wild type, miR-342 deficient and miR-342 Tg mice were subjected for ECAR analysis. Each symbol represents individually tested mouse. *, p<0.05; **, p<0.01; ***, p<0.001; ****, p<0.0001.

Foxp3^DTR^ mice induced for EAE were then treated with DTx at the disease onset, and miR-342 Tg or wild type iTreg cells were adoptively transferred. As shown in Fig 5D, miR-342 Tg Treg cell recipients developed significantly attenuated EAE compared to those of control Treg cell recipients. Furthermore, CNS infiltration of CD4 T cells expressing proinflammatory cytokines was substantially decreased in Tg Treg cell recipients, further demonstrating significantly improved Treg cell functions (Fig 5E). In support, ECAR of Tg Treg cells was significantly diminished even greater than that of wild type Treg cells, while that of Th1 type cells was relatively comparable (Fig 5F). Therefore, these results suggest miR-342-dependent Treg cells’ ability to control inflammatory responses in part by modulating cellular metabolic activity.

## Discussion

We previously demonstrated that miR-342-3p plays an important role in Treg cell suppressive function partly by regulating their metabolic programs ^10^. In this report, we extended the previous study utilizing novel animal models in which miR-342 is targeted or overexpressed in Treg cells and report that miR-342 is critical for Treg cells’ ability to control autoimmune inflammation by modulating glycolytic gene expression. miR-342 expression in Treg cells does not have a major impact on the immune homeostasis under the steady-state conditions, while miR-342 appears to be instrumental for Treg cells’ regulatory functions during inflammation. RNAseq analysis examining the effects of miR-342 deficiency in Treg cells uncovered a striking upregulation of metabolic processes, especially glycolysis. Glycolysis was identified as a top pathway highly enriched in miR-342 deficient Treg cells. Even more striking is that virtually every gene encoding the glycolytic enzymes was substantially upregulated in Treg cells by the lack of miR-342, highlighting miR-342 as a key regulator of cellular glycolysis.

Cell metabolism plays a fundamental role in immune cells, especially regulatory T cells. While Treg cells are known to preferentially utilize oxidative phosphorylation as a main energy source ^33^, there is increasing evidence that Treg cells exploit various metabolic processes to carry out different functions. For instance, Treg cell proliferation and migration requires increased glycolysis, whereas Treg cell suppressive function is wired into oxidative phosphorylation and fatty acid oxidation together with decreased glycolysis ^34^. In support, unconstrained glycolysis in Treg cells could result in diminished Treg cell suppressive function ^35^. Our findings support the notion that regulation of glycolytic pathway is crucial for Treg cell suppressive functions. Indeed, the impact of miR-342 deficiency resulted in defective Treg cell suppressive function, while Treg cell accumulation in the inflamed tissues remained unaffected. Interestingly, diminished oxidative phosphorylation was observed in miR-342 deficient Treg cells. Whether substantially increased glycolysis is responsible for the changes, or alternatively, whether it is mainly due to direct impact of miR-342 deficiency on oxidative phosphorylation remains to be examined.

A single miRNA could target multiple genes ^36^. Our finding that miR-342 deficiency is associated with glycolytic gene upregulation in Treg cells is interesting, especially because every gene involved in glycolytic pathways is found upregulated by miR-342 deficiency. Therefore, miR-342 may serve as a master regulator of cellular glycolysis in Treg cells. The current study also confirms our previous finding that Rictor is targeted by miR-342 ^10^, as its expression is greatly upregulated in miR-342 deficient Treg cells. Unexpectedly, miR-342’s ability to limit Rictor expression was not noticed in Th1 type effector cells. This result strongly suggests that miR-342 may have different targets among different linage CD4 T cells. The target landscape of miR-342 between different effector T cell subsets will require future investigation. miR-342-3p has been shown to directly repress E2F1 expression in human lung cancer cells, from which Myc transcriptional activity is controlled ^12^. While it is possible that miR-342 deficiency may trigger E2F1/Myc-induced cellular glycolysis, we found that E2F1 expression in Treg cells remained unaffected by miR-342 deficiency. Therefore, a direct suppression of E2F1 does not appear to be a mechanism underlying miR-342-dependent Treg cell metabolism/functions. In case of triple negative breast cancer or hepatocellular carcinoma cells, miR-342-3p may target the monocarboxylate transporter 1 (MCT1), which alters lactate and glucose fluxes, further disrupting the metabolic homeostasis, proliferation, and migration of tumor cells ^37^. In support, miR-342-3p expression attenuates tumor cell growth in a xenograft model ^38^. However, glucose uptake measured by 2-NBDG labeling did not reveal notable differences between miR-342 deficient and wild type Treg cells (data not shown). Besides metabolic target genes, miR-342-3p was also shown to regulate NSCLC cell proliferation and apoptosis by targeting Bcl2 ^38^. miR-342-3p/SOCS6 axis regulates protective immunity by increasing inflammatory cytokine and chemokine production^39^.

Methylation status of the CpG island located in the promoter of the host Evl gene may play an important role in miR-342 expression. From analyzing 132 primary B cell lymphoma samples, the Evl locus was found heavily methylated, which was then associated with lower miR-342-3p expression. 5-Aza treatment was able to restore miR-342 and Evl expression by demethylating the promoter, and this was associated with decrease of LC3-II, a marker of autophagy and pro-survival of B lymphoma cells ^40^.

Similarly, epigenetic silencing of the Evl locus was also found in multiple myeloma cells^40^. CpG island methylation-dependent Evl/miR-342 expression was also found during osteogenic differentiation ^41^. miR-342 silencing impairs osteoblast activity and matrix mineralization, whereas its overexpression promotes osteoblast differentiation, directly targeting ATF3, which then inhibits pro-osteogenic differentiation associated gene expression ^41^. Interestingly, relative expression level of pre- and mature miR-342 was not significantly different between Treg and non-Treg (i.e., Th1 type effector) cells. Yet, miR-342-targeted metabolic profiles were evident in Treg but not in Th1 type cells, suggesting that miR-342 may have different target cells between different T cell subsets.

Also important to note is that miR-342 mutant mice reported in this study likely have altered expression of both miR-342-3p and miR-342-5p. Generally, one of the 5’ arm or the 3’ arm of a miRNA precursor is incorporated into the RISC (RNA induced silencing complex) to be functional, and the other arm becomes nonfunctional ^42, 43^, although there is some evidence that both arms could be associated with the RISC and become functional ^44, 45^. Numerous studies using antagomirs or mimics only targeted one of the arms. Therefore, in case both arms are active and functional, targeting both arms of a miRNA vs. the use of antogomirs and/or mimics for one of the arms might have different outcomes. Indeed, miR-342-5p expression could be increased in macrophages within early atherosclerotic lesions in Apoe^-/-^ mice and it directly targets Akt1, which then induces proinflammatory mediator expression (such as Nos2 and Il6) in macrophages via upregulation of miR-155 ^46^. It may be important to examine the targets of each arm in Treg cells.

In sum, our results demonstrate a function of miR-342 that broadly regulates Treg cell metabolism and functions. Given miR-342’s ability to control inflammation and to suppress tumor, exploring the precise mechanisms underlying miR-342-dependent immune regulatory functions may have implications in areas where Treg cells play a key role, including chronic inflammation and tumors.

---

**Supp Figure S1.**
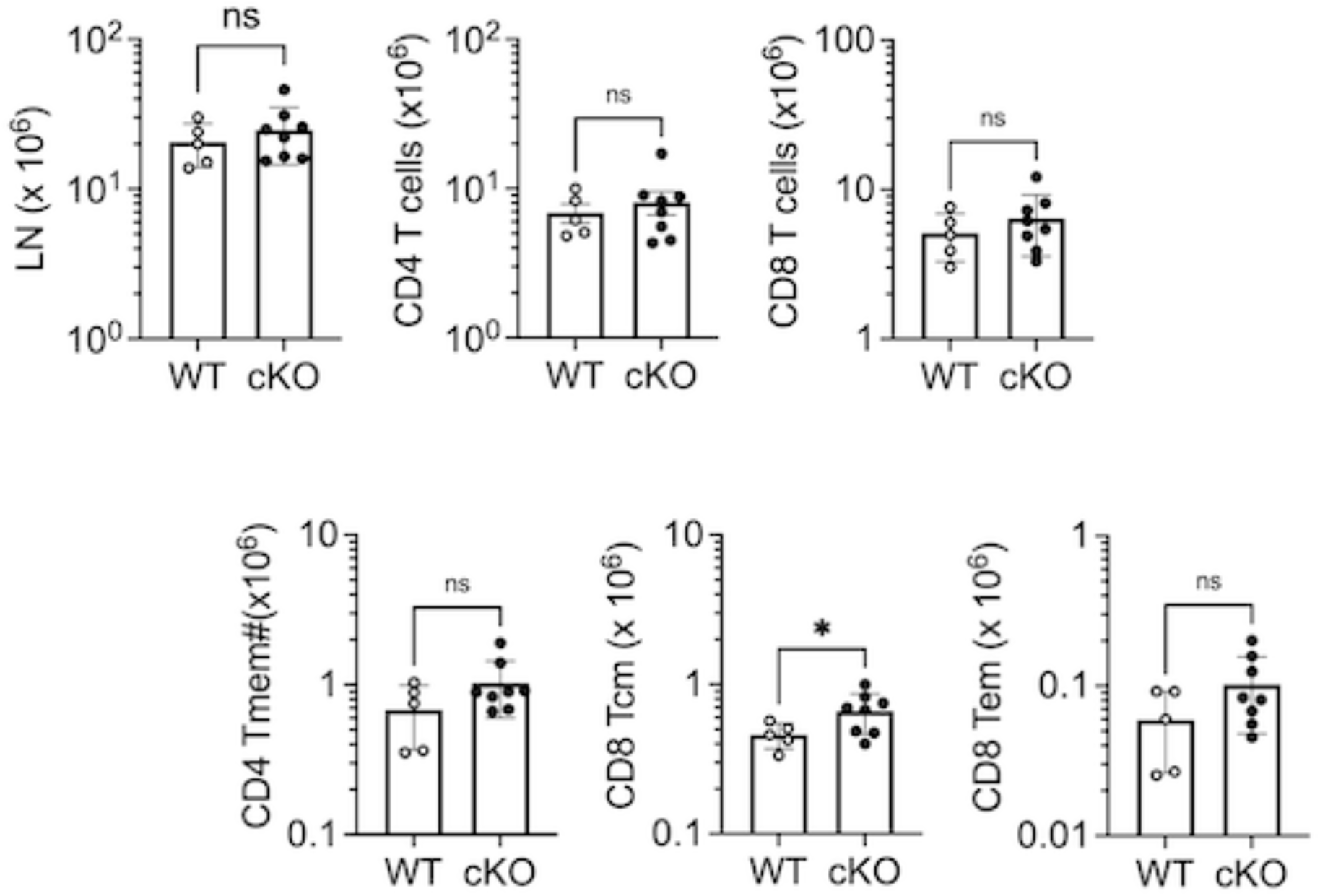
Absolute numbers of lymph node T cell subsets in Treg cell specific miR-342 deficient and wild type mice. Lymph nodes were collected from wild type and miR-342 deficient mice. Absolute numbers of total lymph nodes, CD4, CD8, CD4 memory phenotype, central memory CD8, and effector memory CD8 T cells were enumerated. Each symbol represents individually tested mouse.

**Supp Figure S2.**
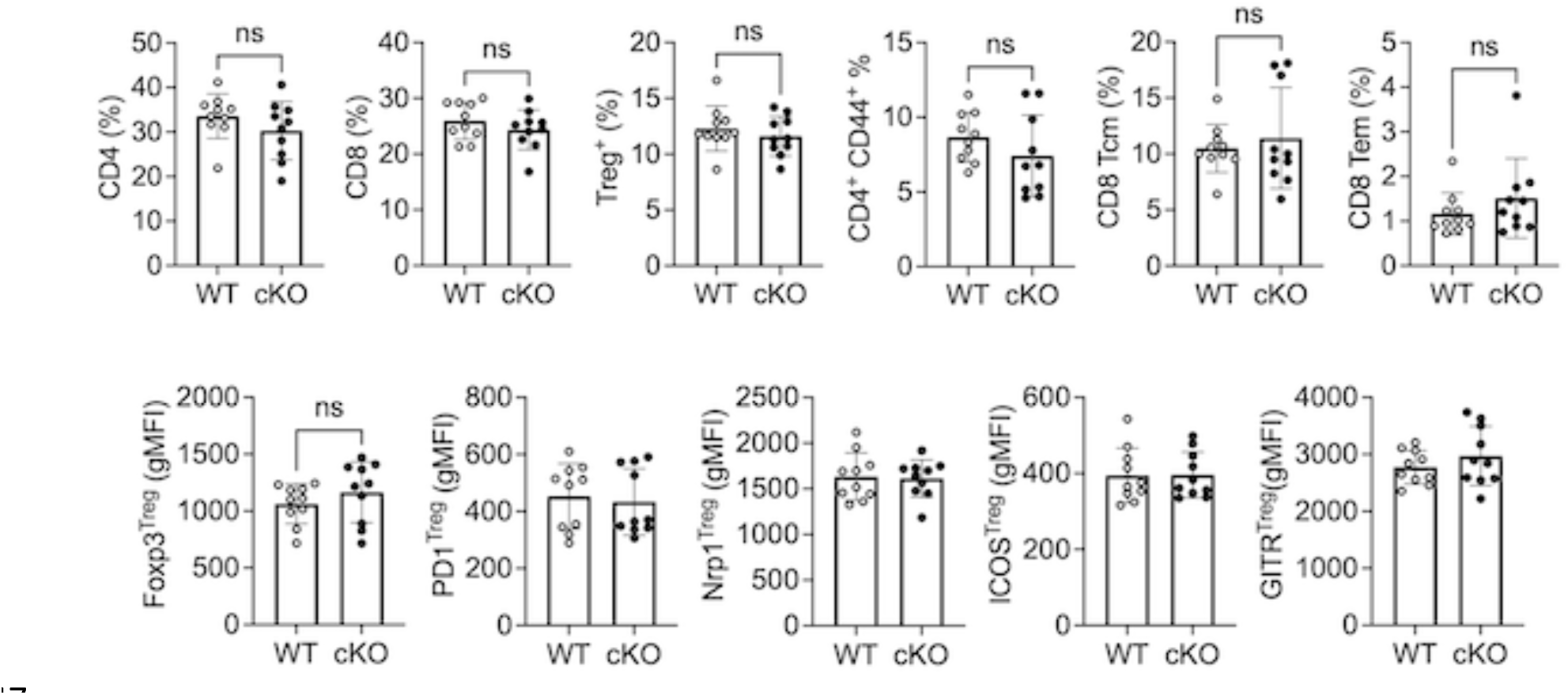
Lymph nodel T cell subsets of T cell specific miR-342 deficient and littermate control mice. Lymph nodes of T^Δ*mir*342^ and control mice were examined for CD4, CD8, Treg, and memory phenotype cells. The data shown are the proportions of each subset. Treg cell expression of Foxp3, PD1, Nrp1, ICOS, and GITR was determined by flow analysis. Each symbol represents individually tested mouse.

**Supp Figure S3.**
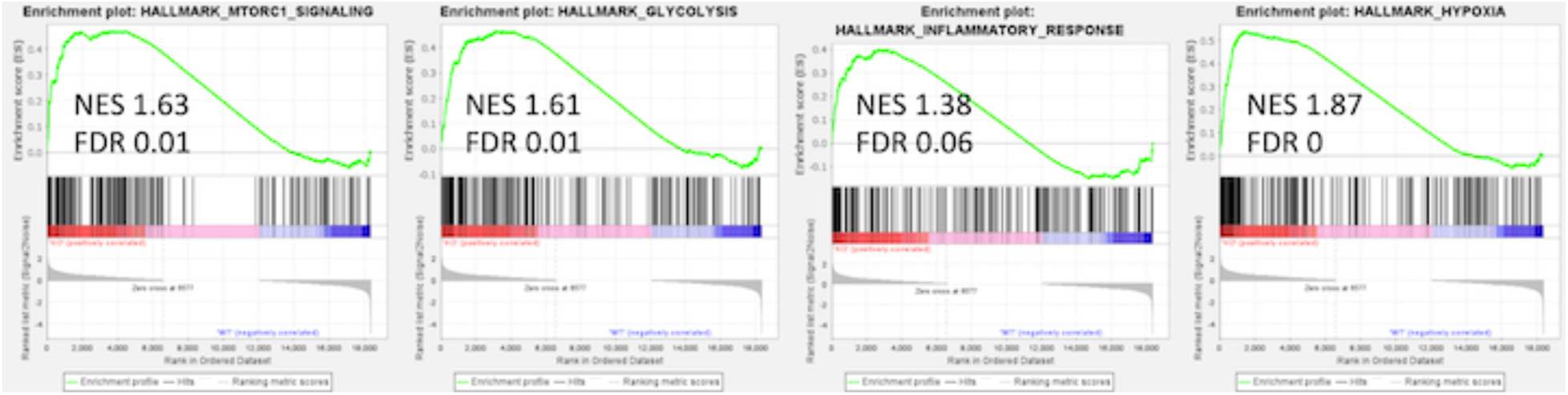
GSEA analysis of Treg^Δ*mir*342^ and control mice. Normalized counts from RNAseq experiments were subjected to Gene Set Eenrichment Analysis (GSEA) based on the hallmark gene sets. The plots shown are top pathways highly enriched in miR-342 deficient Treg cells. The normalized enrichment score (NES) and false discovery rate (FDR) are shown for each gene set.

## References

1. Sakaguchi S, Yamaguchi T, Nomura T, Ono M. Regulatory T Cells and Immune Tolerance. Cell (2008); 133:775–87.

2. Facciabene A, Motz GT, Coukos G. T-regulatory cells: key players in tumor immune escape and angiogenesis. Cancer Res (2012); 72:2162–71. PMC3342842

3. McRitchie BR, Akkaya B. Exhaust the exhausters: Targeting regulatory T cells in the tumor microenvironment. Frontiers in Immunology (2022); 13.

4. Cai Y, Yu X, Hu S, Yu J. A Brief Review on the Mechanisms of miRNA Regulation. Genomics, Proteomics & Bioinformatics (2009); 7:147–54.

5. Friedman RC, Farh KK-H, Burge CB, Bartel DP. Most mammalian mRNAs are conserved targets of microRNAs. Genome research (2009); 19:92–105.

6. Zhou X, Jeker LT, Fife BT, Zhu S, Anderson MS, McManus MT, et al. Selective miRNA disruption in T reg cells leads to uncontrolled autoimmunity. Journal of Experimental Medicine (2008); 205:1983–91.

7. Lu LF, Boldin MP, Chaudhry A, Lin LL, Taganov KD, Hanada T, et al. Function of miR-146a in controlling Treg cell-mediated regulation of Th1 responses. Cell (2010); 142:914–29. PMC3049116

8. Testa U, Pelosi E, Castelli G, Labbaye C. miR-146 and miR-155: Two Key Modulators of Immune Response and Tumor Development. Noncoding RNA (2017); 3. PMC5831915

9. Yao R, Ma YL, Liang W, Li HH, Ma ZJ, Yu X, et al. MicroRNA-155 modulates Treg and Th17 cells differentiation and Th17 cell function by targeting SOCS1. PLoS One (2012); 7:e46082. PMC3473054

10. Kim D, Nguyen QT, Lee J, Lee SH, Janocha A, Kim S, et al. Anti-inflammatory Roles of Glucocorticoids Are Mediated by Foxp3(+) Regulatory T Cells via a miR-342-Dependent Mechanism. Immunity (2020); 53:581–96.e5. PMC7793548

11. Herbst F, Lang TJL, Eckert ESP, Wünsche P, Wurm AA, Kindinger T, et al. The balance between the intronic miR-342 and its host gene Evl determines hematopoietic cell fate decision. Leukemia (2021); 35:2948–63.

12. Tai MC, Kajino T, Nakatochi M, Arima C, Shimada Y, Suzuki M, et al. miR-342-3p regulates MYC transcriptional activity via direct repression of E2F1 in human lung cancer. Carcinogenesis (2015); 36:1464–73.

13. Liu W, Kang L, Han J, Wang Y, Shen C, Yan Z, et al. miR-342-3p suppresses hepatocellular carcinoma proliferation through inhibition of IGF-1R-mediated Warburg effect. Onco Targets Ther (2018); 11:1643–53. PMC5870664

14. Angelin A, Gil-de-Gómez L, Dahiya S, Jiao J, Guo L, Levine MH, et al. Foxp3 Reprograms T Cell Metabolism to Function in Low-Glucose, High-Lactate Environments. Cell Metab (2017); 25:1282–93.e7. PMC5462872

15. Gerriets VA, Kishton RJ, Johnson MO, Cohen S, Siska PJ, Nichols AG, et al. Foxp3 and Toll-like receptor signaling balance T(reg) cell anabolic metabolism for suppression. Nat Immunol (2016); 17:1459–66. PMC5215903

16. Xiao C, Calado DP, Galler G, Thai T-H, Patterson HC, Wang J, et al. MiR-150 Controls B Cell Differentiation by Targeting the Transcription Factor c-Myb. Cell (2007); 131:146–59.

17. Kim D, Le HT, Nguyen QT, Kim S, Lee J, Min B. Cutting Edge: IL-27 Attenuates Autoimmune Neuroinflammation via Regulatory T Cell/Lag3-Dependent but IL-10-Independent Mechanisms In Vivo. J Immunol (2019); 202:1680–5. PMC6401226

18. Dobin A, Davis CA, Schlesinger F, Drenkow J, Zaleski C, Jha S, et al. STAR: ultrafast universal RNA-seq aligner. Bioinformatics (2013); 29:15–21. PMC3530905

19. Liao Y, Smyth GK, Shi W. featureCounts: an efficient general purpose program for assigning sequence reads to genomic features. Bioinformatics (2014); 30:923–30.

20. Love MI, Huber W, Anders S. Moderated estimation of fold change and dispersion for RNA-seq data with DESeq2. Genome Biol (2014); 15:550. PMC4302049

21. Wickham H. Ggplot2: elegant graphics for data analysis. New York: Springer; 2009.

22. Subramanian A, Tamayo P, Mootha VK, Mukherjee S, Ebert BL, Gillette MA, et al. Gene set enrichment analysis: a knowledge-based approach for interpreting genome-wide expression profiles. Proc Natl Acad Sci U S A (2005); 102:15545–50. PMC1239896

23. Mootha VK, Lindgren CM, Eriksson KF, Subramanian A, Sihag S, Lehar J, et al. PGC-1alpha-responsive genes involved in oxidative phosphorylation are coordinately downregulated in human diabetes. Nat Genet (2003); 34:267–73.

24. Zhou Y, Zhou B, Pache L, Chang M, Khodabakhshi AH, Tanaseichuk O, et al. Metascape provides a biologist-oriented resource for the analysis of systems-level datasets. Nat Commun (2019); 10:1523. PMC6447622

25. Foltyn M, Luger A-L, Lorenz NI, Sauer B, Mittelbronn M, Harter PN, et al. The physiological mTOR complex 1 inhibitor DDIT4 mediates therapy resistance in glioblastoma. British Journal of Cancer (2019); 120:481–7.

26. Contreras-Baeza Y, Sandoval PY, Alarcón R, Galaz A, Cortés-Molina F, Alegría K, et al. Monocarboxylate transporter 4 (MCT4) is a high affinity transporter capable of exporting lactate in high-lactate microenvironments. J Biol Chem (2019); 294:20135–47. PMC6937558

27. Parra-Bonilla G, Alvarez DF, Al-Mehdi AB, Alexeyev M, Stevens T. Critical role for lactate dehydrogenase A in aerobic glycolysis that sustains pulmonary microvascular endothelial cell proliferation. Am J Physiol Lung Cell Mol Physiol (2010); 299:L513–22. PMC2957419

28. Roberts DJ, Miyamoto S. Hexokinase II integrates energy metabolism and cellular protection: Akting on mitochondria and TORCing to autophagy. Cell Death Differ (2015); 22:248–57. PMC4291497

29. Kim Y-K, Kim VN. Processing of intronic microRNAs. The EMBO Journal (2007); 26:775–83.

30. Lin SL, Miller JD, Ying SY. Intronic microRNA (miRNA). J Biomed Biotechnol (2006); 2006:26818. PMC1559912

31. Ruby JG, Jan CH, Bartel DP. Intronic microRNA precursors that bypass Drosha processing. Nature (2007); 448:83–6.

32. Kelley K, Chang S-JE, Lin S-L. Mechanism of Repeat-Associated MicroRNAs in Fragile X Syndrome. Neural Plasticity (2012); 2012:104796.

33. Kornberg MD. The immunologic Warburg effect: Evidence and therapeutic opportunities in autoimmunity. Wiley Interdiscip Rev Syst Biol Med (2020); 12:e1486. PMC7507184

34. Kempkes RWM, Joosten I, Koenen HJPM, He X. Metabolic Pathways Involved in Regulatory T Cell Functionality. Frontiers in Immunology (2019); 10.

35. Huynh A, DuPage M, Priyadharshini B, Sage PT, Quiros J, Borges CM, et al. Control of PI(3) kinase in Treg cells maintains homeostasis and lineage stability. Nature Immunology (2015); 16:188–96.

36. Hashimoto Y, Akiyama Y, Yuasa Y. Multiple-to-multiple relationships between microRNAs and target genes in gastric cancer. PLoS One (2013); 8:e62589. PMC3648557

37. Romero-Cordoba SL, Rodriguez-Cuevas S, Bautista-Pina V, Maffuz-Aziz A, D’Ippolito E, Cosentino G, et al. Loss of function of miR-342-3p results in MCT1 over-expression and contributes to oncogenic metabolic reprogramming in triple negative breast cancer. Scientific Reports (2018); 8:12252.

38. Chen Z, Ying J, Shang W, Ding D, Guo M, Wang H. miR-342-3p Regulates the Proliferation and Apoptosis of NSCLC Cells by Targeting BCL-2. Technol Cancer Res Treat (2021); 20:15330338211041193. PMC8445541

39. Fu B, Lin X, Tan S, Zhang R, Xue W, Zhang H, et al. MiR-342 controls Mycobacterium tuberculosis susceptibility by modulating inflammation and cell death. EMBO reports (2021); 22:e52252.

40. Zhang MY, Calin GA, Yuen KS, Jin DY, Chim CS. Epigenetic silencing of miR-342-3p in B cell lymphoma and its impact on autophagy. Clin Epigenetics (2020); 12:150. PMC7574348

41. Han Y, Zhang K, Hong Y, Wang J, Liu Q, Zhang Z, et al. miR-342-3p promotes osteogenic differentiation via targeting ATF3. FEBS Letters (2018); 592:4051–65.

42. Starega-Roslan J, Koscianska E, Kozlowski P, Krzyzosiak WJ. The role of the precursor structure in the biogenesis of microRNA. Cell Mol Life Sci (2011); 68:2859–71. PMC3155042

43. Medley JC, Panzade G, Zinovyeva AY. microRNA strand selection: Unwinding the rules. Wiley Interdiscip Rev RNA (2021); 12:e1627. PMC8047885

44. Ghildiyal M, Xu J, Seitz H, Weng Z, Zamore PD. Sorting of Drosophila small silencing RNAs partitions microRNA* strands into the RNA interference pathway. Rna (2010); 16:43–56. PMC2802036

45. Okamura K, Liu N, Lai EC. Distinct mechanisms for microRNA strand selection by Drosophila Argonautes. Mol Cell (2009); 36:431–44. PMC2785079

46. Wei Y, Nazari-Jahantigh M, Chan L, Zhu M, Heyll K, Corbalán-Campos J, et al. The *microRNA-342-5p* Fosters Inflammatory Macrophage Activation Through an Akt1- and *microRNA-155*&#x2013;Dependent Pathway During Atherosclerosis. Circulation (2013); 127:1609–19.

